# A label-free approach to detect ligand binding to cell surface proteins in real time

**DOI:** 10.1101/251629

**Authors:** Verena Burtscher, Matej Hotka, Yang Li, Michael Freissmuth, Walter Sandtner

## Abstract

Electrophysiological recordings allow for monitoring the operation of proteins with high temporal resolution down to the single molecule level. This technique has been exploited to track either ion flow arising from channel opening or the synchronized movement of charged residues and/or ions within the membrane electric field. Here, we describe a novel type of current by using the serotonin transporter (SERT) as a model. We examined transient currents elicited on rapid application of specific SERT inhibitors. Our analysis shows that these currents originate from ligand binding and not from a conformational change. The Gouy-Chapman model predicts that a ligand-induced elimination/neutralization of surface charge must produce a displacement current and related apparent changes in membrane capacitance. Here we verified these predictions with SERT. Our observations demonstrate that ligand binding to a protein can be monitored in real time and in a label-free manner by recording the membrane capacitance.

## Introduction

Voltage-clamp recordings have been used for the past 70 years to assess currents through membranes, which are either resistive (i.e. ion flux through ion channels, Hodgkin & Huxley 1952) or which arise from synchronized movement of charged residues or ions within the membrane electric field (i.e. gating currents, Armstrong & Bezanilla 1973). The two hallmarks of electrophysiological recordings are high sensitivity and high temporal resolution: it is technically feasible to monitor currents through single molecules and to resolve conformational transitions, which occur on the microsecond scale. In addition, voltage-clamp recordings also allow for determining the cell membrane capacitance. By this approach, it has been possible to detect changes in membrane area that result from the fusion of a single vesicle with the plasma membrane (Neher and Marty, 1982; Fernandez et al., 1984; von Gersdorff and Matthews, 1994). However, a change in membrane area is not the only factor, which can affect capacitance. Several studies – most of them conducted in cells that overexpressed a membrane protein – demonstrated the contribution of the same to the membrane capacitance: the effect can be accounted for by mobile charges within the protein. These add to the total charge required to recharge the membrane upon prior voltage change. Voltage-gated channels were the first membrane proteins, on which capacitance measurements were conducted: the total membrane capacitance of a cell expressing these proteins was shown to be comprised of a voltage-independent component, which was ascribed to the membrane, and of a voltage-dependent component, which was attributed to mobile charges within the voltage sensors of the channel (Bean and Rios, 1989). However, capacitance measurements also allow for extracting information on the conformational switch associated with binding of ions and/or substrate to transporters: it has, for instance, been possible to infer the conformational transition associated with phosphorylation of the Na^+^/K^+^-pump from the recorded change in capacitance (Lu et al., 1995). Similarly, binding of the co-substrate Cl^-^ to the inward facing conformation of the γ-aminobutyric acid transporter-1 (GAT1/SLC6A1) caused a very rapid reduction in the recorded capacitance, which was proposed to reflect the suppression of a charge moving reaction induced by binding of Na^+^ to the outward facing conformation of GAT1.

In the present study, we conducted current and capacitance measurements in HEK293 cells expressing human SERT. Like GAT1, SERT is a member of the solute carrier 6 (SCL6) family. The transport cycle of SERT is understood in considerable detail (Schicker et al., 2012; Sandtner et al., 2014), in particular the nature of substrate-induced currents, electrogenic steps associated with binding and release of co-substrate ions and decision points in the transport cycle (Hasenhuetl et al., 2016; Kern et al., 2017). In addition SERT, in contrast to GAT1, has a very rich pharmacology (Sitte and Freissmuth, 2015), which allows for probing the actions of inhibitors (Hasenhuetl et al., 2015), atypical inhibitors (Sandtner et al., 2016), substrate and releasers (Sandtner et al., 2014; Kern et al., 2017), and atypical substrates/partial releasers (Bhat et al., 2017) by electrophysiological recordings. Here we explored changes in apparent membrane capacitance resulting from ligand binding to SERT. A reduction in membrane capacitance was seen regardless of whether the ligand was the cognate substrate or an inhibitor of SERT, e.g. cocaine. The Gouy-Chapman model (Gouy, 1909; Chapman, 1913) predicts that charge neutralisation must result in the apparent reduction of the membrane capacitance and in ligand-induced displacement currents. We verified these predictions, by analysing both the ligand-induced currents and the related changes in membrane capacitance.

## Results

### Application of cocaine to HEK293 cells stably expressing SERT induces an inwardly directed peak current

When measured in the whole cell configuration, rapid application of 5-HT to SERT gives rise to an initial peak current, which is followed by a steady state current (Schicker et al., 2012). The peak current is caused by a synchronized conformational rearrangement that carries 5-HT through the membrane (Hasenhuetl et al., 2016). The steady-state current on the other hand is contingent on the progression of SERT through the transport cycle. Raising intracellular Na^+^ impedes this cycle and eliminates the steady-state current (Hasenhuetl et al., 2015). This allows for studying the peak current in isolation as shown in the upper panel of Fig. 1a, which depicts a representative trace elicited by 30 μM 5-HT in the presence of 152 mM intracellular Na^+^.

**Figure 1.**
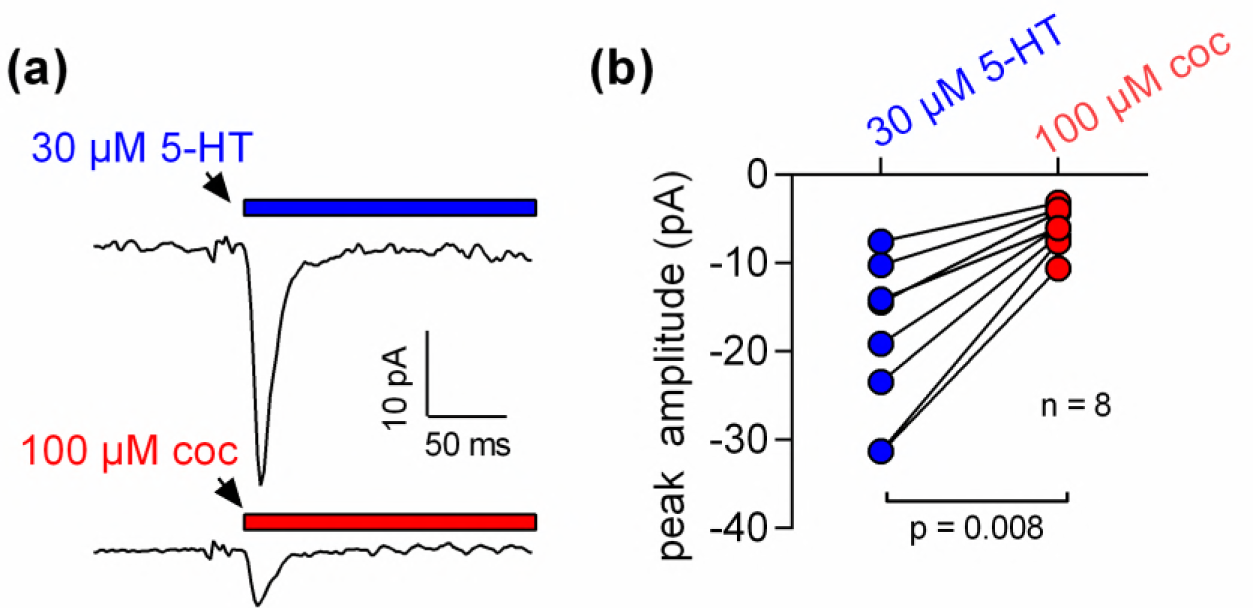
5-HT and cocaine give rise to inwardly directed peak currents when applied to HEK293 cells stably expressing SERT. (a) Representative currents from the same cell evoked by rapid application of 30 μM 5-HT (upper panel) or 100 μM cocaine (lower panel), respectively. The currents were recorded at 0 mV and in the presence of 152 mM intracellular Na^+^ (b) Comparison of the amplitudes of the 5-HT (blue circles) and the cocaine induced peak current (red circles). The lines connect measurements from the same cell: 5-HT: −18.8 ± 9.1 pA; cocaine: −6.0 ± 2.4 pA; n=8; p=0.008, Wilcoxon test.

We applied 100 μM cocaine to a cell expressing SERT to explore, if inhibitors of SERT also caused conformational rearrangements detectable by electrophysiological recordings. The recording in the lower panel of Fig. 1a shows a cocaine-induced current, which was smaller than that elicited by 5-HT and which was absent in control cells (data not shown): In paired measurement, where cells were challenged with both, 30 μM 5-HT and 100 μM cocaine (Fig. 1b), we observed a constant amplitude ratio (5-HT: cocaine ∼ 4:1).

### The amplitude and the kinetics of the cocaine-induced current are concentration dependent

The K_D_ for cocaine binding to SERT has been estimated by radioligand binding assays, by the concentration dependence of its inhibitory action on substrate uptake and by electrophysiological means (Hasenhuetl et al., 2015). The affinity estimates provided by these measurements range from 100 nM to 3 μM (Han and Gu, 2006; Hasenhuetl et al., 2015). These concentrations are much lower than the 100 μM used in the above experiments. Accordingly, we measured the current in response to a wide range of cocaine concentrations (Fig. 2a). It is evident from these recordings that an increase in cocaine concentration was associated with larger amplitudes and accelerated peak current decays. The time courses of the decays were adequately fit by a monoexponential function. In Fig 2b, we plotted the rate estimates by the fits as a function of the applied cocaine concentration. The rates increased linearly over the tested concentration range. At concentrations below 10 μM, cocaine failed to evoke detectable currents.

**Figure 2.**
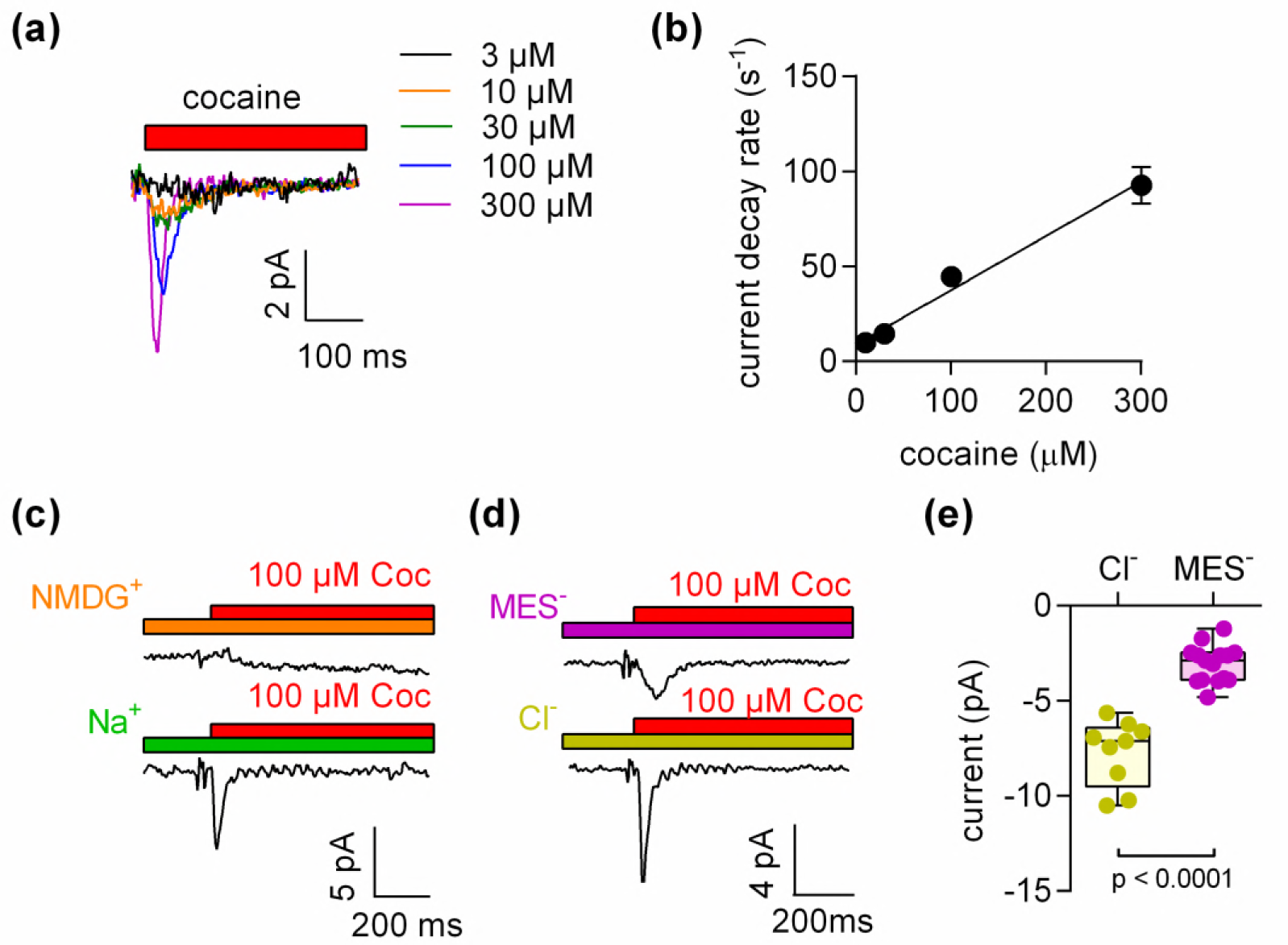
The amplitude of the cocaine-induced current and the time-course of its decay are concentration dependent. (a) Representative currents elicited by 3 μM (black), 10 μM (orange), 30 μM (green), 100 μM (blue) and 300 μM (magenta) cocaine, respectively. The depicted currents were recorded from the same cell. (b) Rate of current decay as a function of cocaine concentration. The line is a linear fit to the data points (slope: 2.9*10^5^ ± 0.3*10^5^ Mol^-1^s^-1^; n=7). Data are mean ± SD. (c) In the absence of Na^+^, 100 μM cocaine failed to induce a peak current. Shown are representative traces in the presence of 150 mM extracellular NMDG^+^ (upper panel) and 150 mM extracellular Na^+^ (lower panel), respectively. The depicted traces were recorded from the same cell. The absence of the cocaine-induced current in the presence of NMDG^+^ was confirmed in 7 independent experiments. (d) Representative currents in the presence of 150 mM extracellular MES^-^ (upper panel) and 150 mM extracellular Cl^-^ (lower panel), respectively. The amplitude of the cocaine-induced current peak was −7.7 ± 1.7 pA (n=9) and −3.1 ± 1.0 pA (n=15) in the presence and absence of Cl^-^, respectively. (p<0.0001; Mann-Whitney U-test)

### The cocaine-induced current requires the presence of extracellular Na^+^

Binding of competitive inhibitors to SERT is dependent on Na^+^ (Korkhov et al., 2006). Therefore, we measured the cocaine peak in the absence of extracellular Na^+^: in the absence of extracellular Na^+^, 100 μM cocaine failed to elicit a current (Fig. 2c). In contrast, when Cl^-^ was removed from the extracellular solution, the cocaine-induced currents were present, but featured smaller amplitudes (Fig. 2d,e).

### The amplitude of the cocaine-induced current depends on voltage and is larger at positive potentials

We measured the voltage dependence of the cocaine-induced current by relying on the protocol depicted in the left panel of Fig. 3a: the membrane was clamped for 1 minute to a constant voltage ranging from −50 to +30 mV. During each of this successive voltage clamps, 100 μM cocaine was applied for 5 seconds and subsequently removed to generate a family of currents (right hand panel of Fig. 3a). Excessive noise precluded analysis above and below this voltage range. It is evident from Fig. 3a that the cocaine peak became larger at positive potentials. The current-voltage relation was examined by averaging 9 independent experiments after normalizing the current amplitudes to the current recorded at +30 mV: this current-voltage relation was linear over the explored voltage range and had a negative slope.

### The absence of Cl^-^ prevents conformational change in SERT and renders the 5-HT induced peak current similar to the current induced by cocaine

It has long been known that extracellular Cl^-^ is required for SERT-mediated substrate uptake (Lingjærde, 1971; Nelson and Rudnick, 1982). However, the presence of Cl^-^ is not required for 5-HT binding to SERT (Nelson and Rudnick, 1982). In fact, the absence of Cl^-^ precludes the conversion of the substrate-loaded outward-facing transporter to the inward-facing conformation. Thus, in the following experiments, we removed Cl^-^ from the extracellular solution to separate the event of 5-HT binding from the subsequent conformational change.

**Figure 3.**
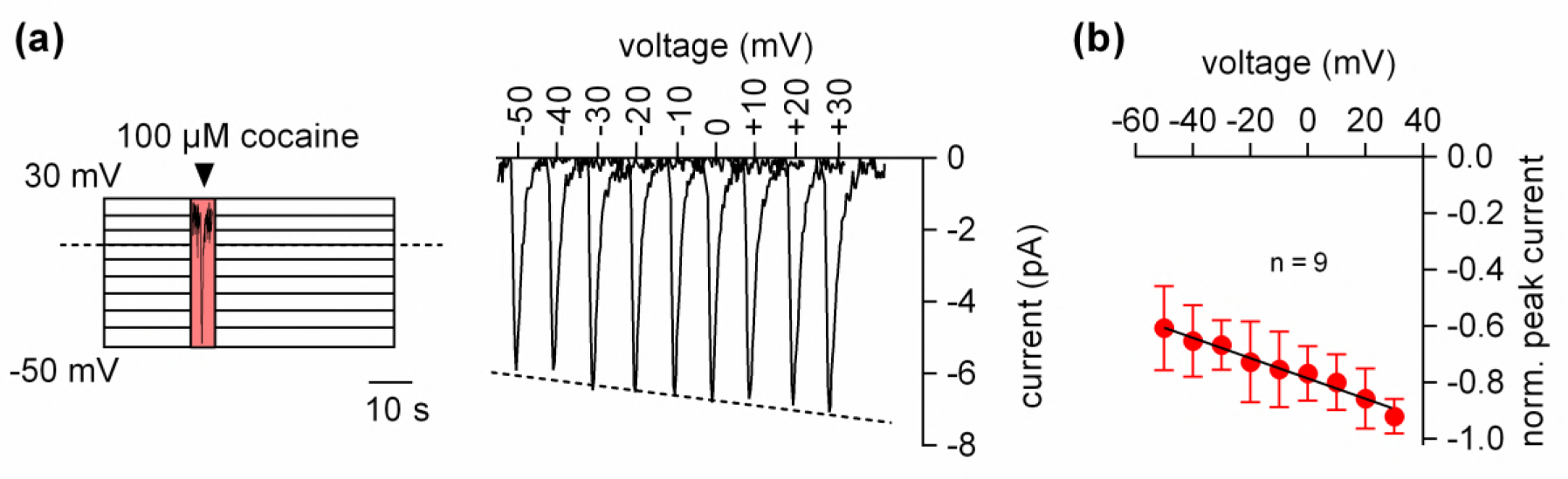
The amplitude of the cocaine peak increases at positive potentials. (a) The scheme illustrates the protocol employed to measure the voltage dependence of the cocaine induced peak current: cells were clamped to voltages between −50 and +30 mV for 60 seconds, respectively. At each potential 100 μM cocaine was applied for 5 seconds and subsequently removed. Shown are representative currents elicited by the protocol. (b) Peak currents induced by 100 μM cocaine were normalized to the current at +30 mV (n= 9). Data are mean ± SD. The black line is a linear fit to the data points (slope= −3.5*10^-3^ ± 5.3*10^-4^/mV).

Removal of Cl^-^ reduced the peak amplitude and the area under the peak (see Fig. 4a for representative current traces and Fig. 4b for the comparison of 20 independent experiments with paired recordings done in the presence and absence of Cl^-^). We also determined the current-voltage relation in the presence (Fig. 4c,d) and absence of Cl^-^ (Fig. 4,e,f): by contrast with the positive slope seen in the presence of Cl^-^ (Fig. 4c,d), the slope was negative in the absence of Cl^-^ (Fig. 4e, f). These observations indicated that the absence of Cl^-^ unmasked an event preceding the substrate-induced conformational change: this initial event had a small current amplitude and a negative slope in the current-voltage relationship.

**Figure 4.**
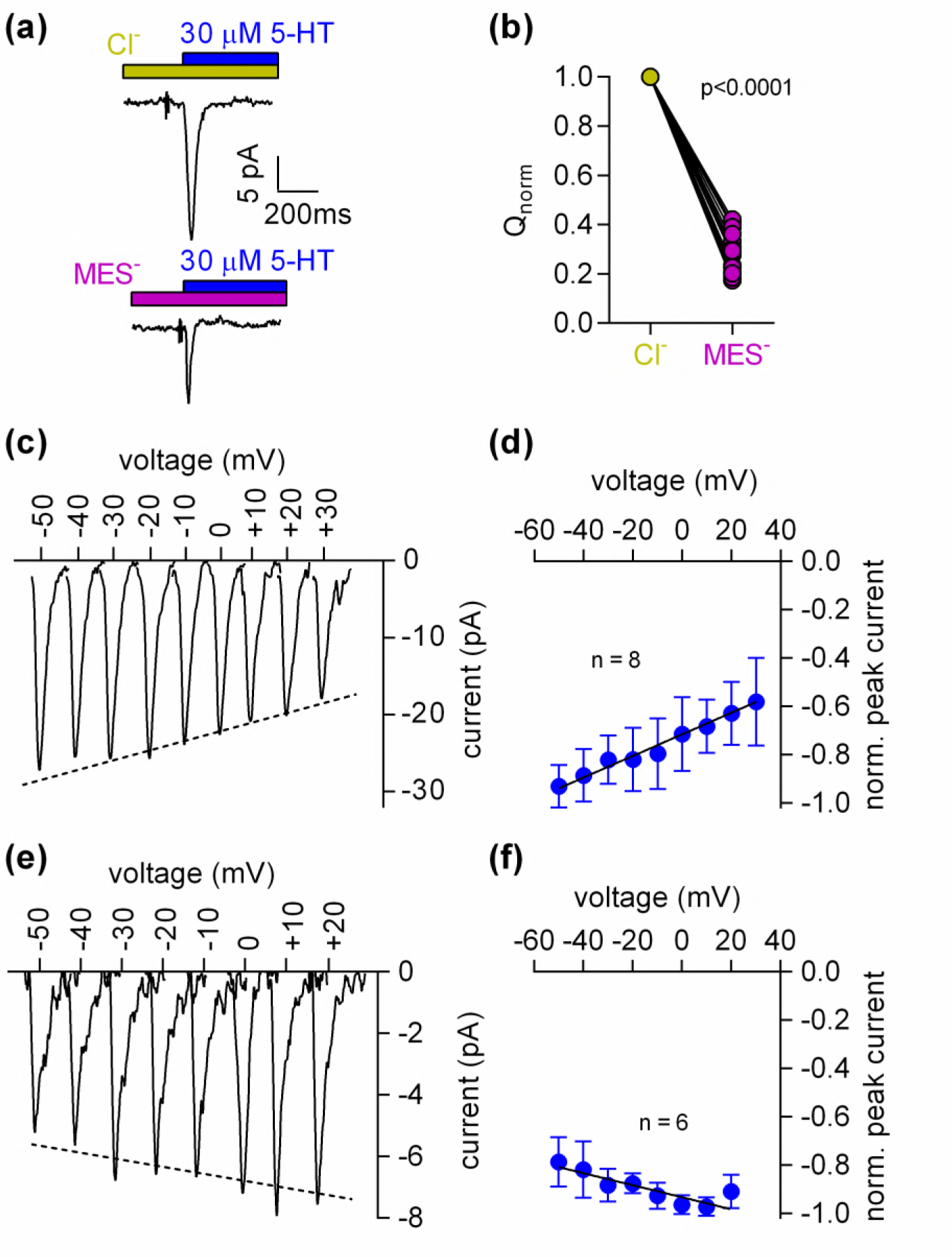
Cl^-^ removal diminishes the amplitude of the 5-HT induced current and reverts the slope of the voltage dependence of the peak current (a) Representative traces of currents evoked by 30 μM 5-HT in the presence of 150 mM extracellular Cl^-^ (upper panel) or 150 mM MES^-^ (lower panel) from the same cell. The cell was held at 0 mV. (b) Comparison of the normalized charge (Q) carried by the 5-HT-induced current in the presence and absence of Cl^-^. Connecting lines indicate measurements from the same cell. After removal of Cl^-^ the remaining fraction of charge was 0.28 ± 0.075 (n=20). The charge before and after Cl^-^ removal was significantly different (p=0.0001; Wilcoxon test) (c) Representative peak currents induced by 30 μM 5-HT recorded at potentials ranging from −50 to +30 mV in the presence of 150 mM extracellular Cl^-^. (d) The amplitude of the 5-HT induced peak current measured in the presence of Cl^-^ as a function of voltage. The peak current amplitudes evoked by 30 μM 5-HT were normalized to the current recorded at −50 mV (n=8). Data are mean ± SD. The line is a linear fit to the data-points (slope= 4.5*10^-3^ ± 5.4*10^-4^/mV). (e) Representative peak currents evoked by 30 μM 5-HT in the absence of Cl^-^ recorded at potentials ranging from −50 to +20 mV. (e) Normalized 5-HT induced peak currents recorded in the absence of Cl-. The peak current amplitudes were normalized to the current recorded at +10 mV (n=6). Data are mean ± SD. The line is a fit to the data points (slope= −2.5*10^-3^ ± 4.9*10^-4^/mV)

### The Gouy-Chapman model predicts displacement currents upon ligand binding to membrane proteins

It is conceivable that binding of cocaine or of 5-HT to SERT occurs via an induced fit, i.e. local rearrangements are required within the vicinity of the binding pocket to accommodate the compound. These rearrangements can be expected to force charged residues of SERT to change their position within the membrane electric field. However, it is difficult to reconcile this with the negative slope in the current-voltage relation of both, the cocaine- and the 5-HT-induced peak current in the absence of Cl^-^. The negative slope in the current-voltage relation was inconsistent with the conjecture that the cocaine-induced peak current originated from charges moving in the membrane electric field. The rules of electrostatics dictate that inwardly directed currents must increase at negative voltages, if they are produced by charges moving in response to a conformational change. This is also evident from the current-voltage relation for the 5-HT induced peak current in the presence of chloride (Fig. 4c & 4d). This current reflects the conformational rearrangement that carries substrate and co-substrates through the membrane (Hasenhuetl et al., 2016). Accordingly, the corresponding current-voltage relation had a positive slope (Fig. 4d).

Therefore, an alternative explanation is more plausible, if it can relate these currents to ligand binding without the need to invoke conformational changes: the Gouy-Chapman model provides a mathematical description of a simplified biological membrane (see scheme in Fig. 5a) and allows for calculating the potentials at the inner and outer surface of the membrane. These potentials depend on the voltage difference between the intra- and extracellular bulk solution, the ionic composition of these solutions and the number of charges at the inner and outer membrane surfaces. In biological membranes, the charges at the surfaces are comprised of the phospholipid head groups and the solvent-accessible acidic or basic residues of membrane proteins. The number of charges at the surfaces is an important determinant of the transmembrane potential (V_trans_), which represents the difference between the potentials at the opposite surfaces. In our stable cell line SERT is expressed at high levels; the number of SERT molecules was estimated in saturation binding experiments of [^3^H]imipramine and amounted to 20 ± 4.2 × 10^6^ transporters/cell. At this expression level, the solvent accessible charge on SERT is expected to contribute significantly to the total surface charge. Assuming one surface charge per transporter we calculated a surface charge density of about 0.00011 C/m^2^. In comparison, the total surface charge density on the outer surface of untransfected HEK293 cells has previously been determined by an experimental approach and amounted to 0.005 C/m^2^ (Zhang et al., 2001). This implies that in the SERT expressing cell line used, up to 2.5 % of the outer surface charge is likely attributable to SERT.

**Figure 5.**
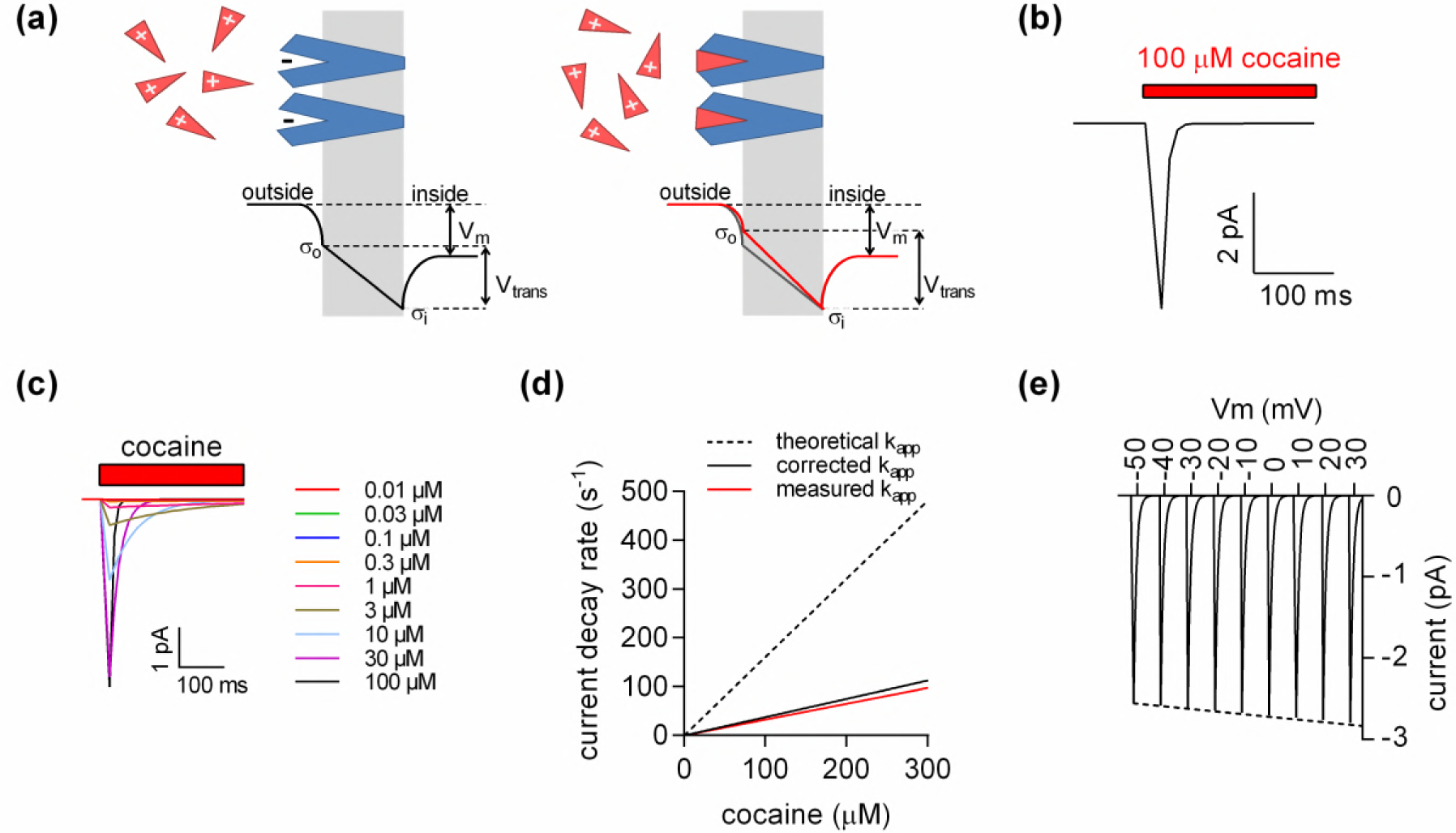
A model for ligand induced surface charge elimination. (a) Scheme of a membrane (grey slab) with embedded transporters (in blue) and ligands (red triangles). The ligand is positively charged and the binding site on the protein comprises a negative charge (left panel). Upon binding the, ligand neutralizes the charge on the protein (right panel). The lines in the left (in black) and right panel (in red) indicate voltage profiles across the membrane. Binding of the ligand to the negative surface charge on the protein renders the outer surface charge potential more positive. This produces a change in transmembrane voltage V_tra_ns (ΔV_tran_s). (b) Displacement current predicted by the Gouy-Chapman model. The instantaneous current calculates as: i(t)=C_m_ · dv(t) · d(t)^-1^, where C_m_ is the capacitance of the membrane and v(t)= V_trans_ · (1-e^-kapp*t^); k_app_ is the apparent rate of cocaine association (k_app_=k_on_ · [cocaine]+k_off_). Shown is a simulated current evoked by application of 100 μM cocaine. (c) Simulated currents at the indicated cocaine concentrations. (d) Predicted rates of the current decays of the cocaine peaks as a function of the cocaine concentration (dashed line). The solid red line in the plot indicates measured rates from figure 2b. The black solid line indicates corrected rates (see supplement, section 2) (e) Simulated voltage dependence of the cocaine peak. The current-voltage relation has a negative slope (slope= −1.1*10^-3^ ± 1.4*10^-5^/mV).

We utilized the Gouy-Chapman model to test the following hypothesis (see also scheme in 5a): ligands of SERT such as 5-HT or cocaine are positively charged. The corresponding binding site on SERT on the other hand holds at least one negative charge (D98, Barker et al. 1999; Celik et al. 2008). We posited that the latter is a surface charge that can be neutralized/eliminated by the ligand. This idea was incorporated into the model, which predicted a change in V_trans_ of about 10 mV upon ligand binding. Because, the cell membrane represents a capacitor, a change in transmembrane voltage must result in a displacement current. In figure 5b, we simulated displacement currents based on the voltage estimates provided by the model. The simulated currents are inwardly directed with amplitudes in the low picoampere range and they thus match the observed currents.

In Figure 5c, we modeled the concentration dependence of the cocaine-induced peak: it is implicit to our hypothesis that the time course of ligand-induced change in V_trans_ must coincide with the time course of cocaine binding. We recently determined the association rate (k_on_) and dissociation rate (k_off_) of cocaine for SERT by an electrophysiological approach (Hasenhuetl et al., 2015). These rates were used to calculate the apparent association rate (k_app_, dashed line in Fig. 5d). At low cocaine concentrations, the simulated currents compared favorably with the observed. However, at higher cocaine concentrations the predictions deviated from the measured k_app_ (red solid line in Fig. 5d). We attribute this discrepancy to the fact that, at concentrations exceeding 30 μM, the solution exchange by our application device (-20 s^1^) becomes rate-limiting; for technical reasons, the diffusion-limited association-rate for cocaine is therefore currently inaccessible to an experimental determination. We applied a correction for the finite solution exchange rate (see supplement, section 2), which predicts the corrected k_app_ (black solid line).

We also calculated the current-voltage relation for the displacement current (Fig. 5e): Consistent with our observations, the synthetic data predicts larger currents at positive potentials. The hypothesis that ligand binding results in charge neutralization and generation of a displacement current therefore provides a parsimonious explanation for the negative slope of the observed current-voltage relation.

### Application of cocaine to HEK293 cells expressing SERT decreases the apparent membrane capacitance

The Gouy-Chapman model can be used to calculate the change in apparent membrane capacitance resulting from the ligand-induced reduction in extracellular surface charge (Fig. 6a). This prediction was verified. Figure 6b shows a representative recording of the membrane capacitance upon application and subsequent removal of 100 μM cocaine to HEK293 cells expressing SERT. This effect of cocaine was absent in control cells (lower panel). The reduction in membrane capacitance by cocaine amounted to approx. 500 fF, which was in good agreement with the prediction (Fig.6c). Figure 6d shows the concentrations dependence of the cocaine-induced decrease in apparent membrane capacitance. These data were adequately fit by a saturation hyperbola, which provided an estimate for the affinity of cocaine to SERT (EC_50_ = 156 ± 41 nM) (Fig.6e). This estimate is in line the published K_D_ of cocaine (Hasenhuetl et al., 2015).

**Figure 6.**
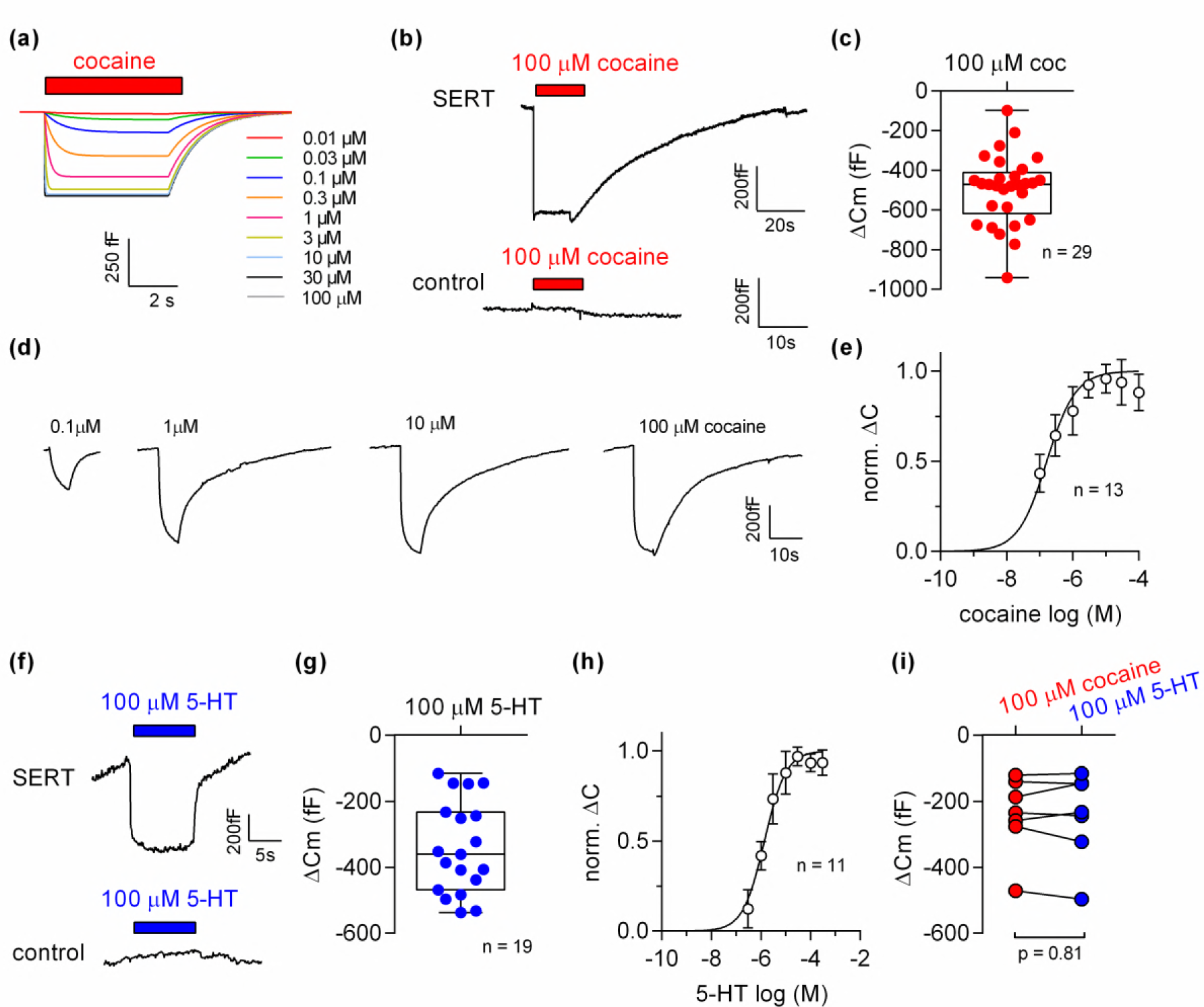
Cocaine and 5-HT binding to SERT results in a reduction of apparent membrane capacitance. (a) Predicted change in membrane capacitance by binding of cocaine by the Gouy-Chapman model. The traces are the simulated response to the indicated cocaine concentrations. (b) Representative change in capacitance recorded in the presence of 100 μM cocaine in a SERT expressing cell (top) and in a control cell (bottom). (c) Plot of the change (ΔCm) induced by 100 μM cocaine (n = 29; ΔCm = −495 ± 175 fF). (d) Representative traces of the cocaine-induced apparent reduction in capacitance at the indicated concentrations. The recordings are from the same cell. (e) Concentration-response curve for the cocaine-induced change in membrane capacitance (n = 13), which was normalized to the maximal cocaine-induced ACm. Data are mean ± SD. The solid line was generated by fitting the data to a rectangular hyperbola yielding an EC_50_= 164 ± 41 nM. (f) Representative change in capacitance elicited by 100 μM 5-HT in a SERT expressing cell and in a control cell. (g) Plot of the change (ACm) induced by 100 μM 5-HT (n = 19, ΔCm = −340 ± 140 fF). (h) Concentration-response curve for the 5-HT induced change in membrane capacitance (n = 11), which was normalized to the change in capacitance at 10 μM cocaine. Data are mean ± SD. The solid line was generated by fitting the data to a rectangular hyperbola yielding an EC_50_= 1.4 ± 0.1 μM. (i) Comparison of the response to 100 μM cocaine and 100 μM 5-HT, respectively (n = 8). The lines connect measured values from the same cell. The data are not significantly different (p = 0.81; Wilcoxon test).

We also explored the effect of 100 μM 5-HT in presence of external Cl^-^ on the apparent membrane capacitance. Application of 5-HT reduced the membrane capacitance in cells expressing SERT (upper panel in Fig. 6f) but not in control cells (lower panel in Fig. 6f). Upon removal of 5-HT, the membrane capacitance relaxed to the initial level in cells expressing SERT (upper panel in Fig. 6f). We applied increasing concentrations of 5-HT to determine the concentration-response curve for the 5-HT-induced apparent reduction in membrane capacitance (Fig. 6g). The resulting saturation hyperbola provided an estimate for the apparent affinity of 5-HT to SERT (EC_50_ = 1.4 ± 0.1 μM, Fig.6h). We also compared the apparent reduction of capacitance elicited by 100 μM cocaine and 100 μM 5-HT. Because 100 μM is a saturating concentration for both ligands, the capacitance change was predicted to be equivalent provided that both compounds neutralized the same charge. This was the case: the magnitude of the reduction was comparable in paired recordings (Fig. 6i).

### The voltage dependence of the cocaine-induced capacitance change is predicted by the Gouy-Chapman model

The voltage dependence of the cocaine-induced capacitance change was analyzed using the protocol outlined in Fig. 7a. We first measured the membrane capacitance in the absence of cocaine at −50 mV, 0 mV and +50 mV and applied 100 μM cocaine thereafter. Consistent with the results shown in Figure 6a application of cocaine reduced the membrane capacitance in SERT expressing cells (left panel in Fig. 7a) but not in control cells (right panel in Fig. 7a). We also simulated the recordings in SERT expressing cells using the Gouy-Chapman model: these synthetic data recapitulated the capacitance recordings (cf. left panel in Fig. 7a and Fig. 7b). The capacitance recordings from 6 independent experiments are summarized in Fig. 7c. The black and the grey dashed lines represent the linear regression through the actual recordings and the changes predicted the Gouy-Chapman model, respectively. It is evident that (i) the cocaine-induced decrease in capacitance depends on the voltage in a linear manner and (ii) that the experimental observations and the synthetic data are in reasonable agreement. However, there is one notable difference: in the absence of cocaine, changes in voltage were followed by a slow change in capacitance in SERT expressing cells, in particular when the prior voltage had been +50 mV (inset Fig.7a). The model does not account for this feature. Nevertheless, this slow change in capacitance is specific to SERT-expressing cells, because it is absent in control cells and inhibited by cocaine. We surmise that the transient change in capacitance reflects entry into / return from a state of SERT, which is occupied at very positive potentials.

**Figure 7.**
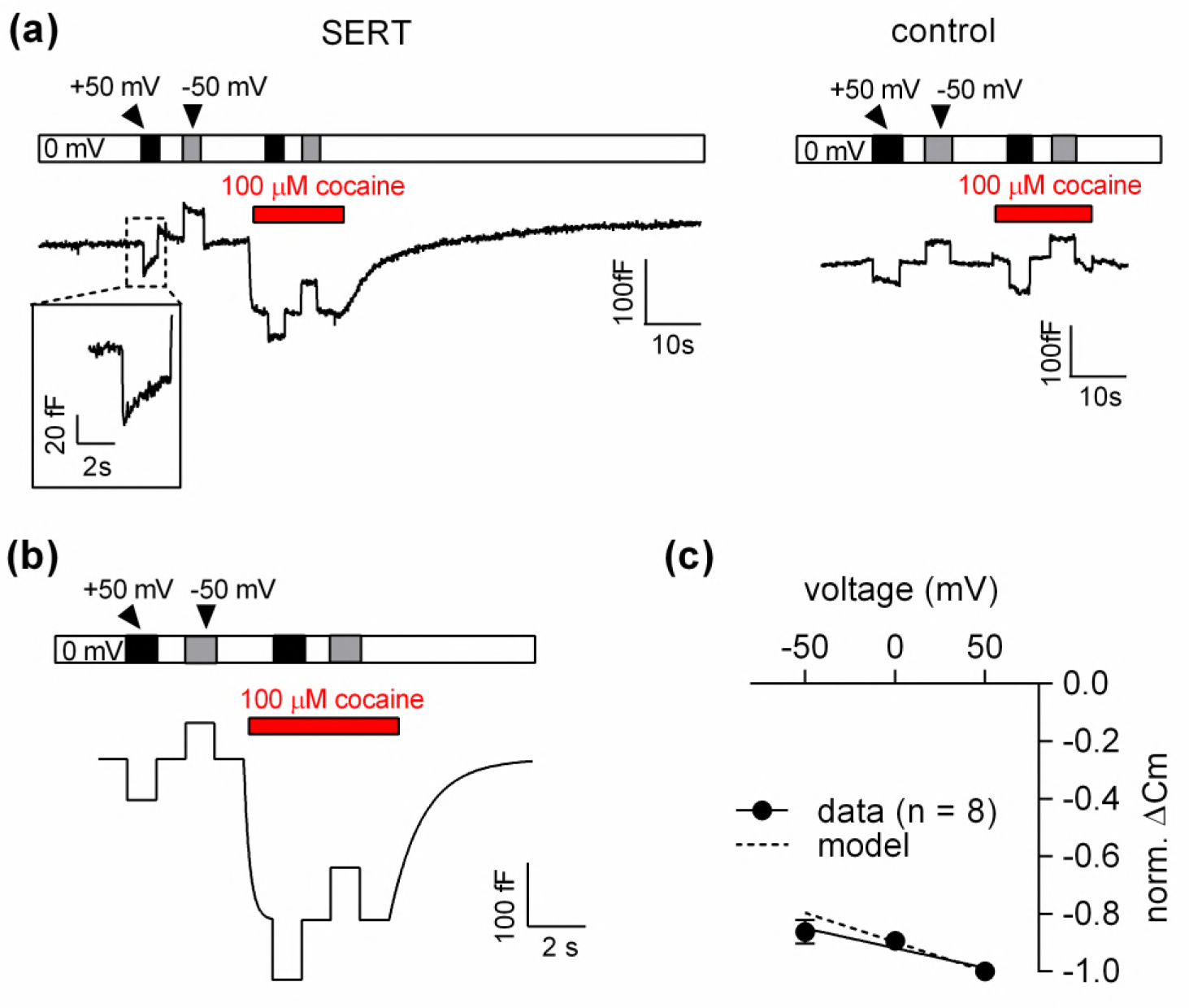
Voltage dependence of the reduction in apparent membrane capacitance by cocaine. (a) The capacitance was recorded in a SERT expressing cell (left panel) and in a control cell (right panel). The holding potential was changed to +50 or −50 mV as shown by the bar in the absence and presence of 100 μM cocaine. (b) The experiment in the left panel (a) was simulated by a model based on the Gouy-Chapman equation. (c) The capacitance was recorded as outlined in Fig. 6b −50 mV, 0 mV and 50 mV and the cocaine-induced change was normalized by setting the amplitude of the capacitance change at +50 mV to −1. Data are means ± SD (n= 6). The solid line was drawn by linear regressions (slope= −1.4*10^-3^ ± 1.7*10^-4^/mV). The dashed line indicates the voltage dependence of the cocaine-induced capacitance change predicted by the Gouy-Chapman model.

### The cocaine-induced current peak carried by SERT exemplifies a more general phenomenon

In the Gouy-Chapman model no reference is made to specific proteins or ligands. In this context, SERT is treated as an entity, which contributes to the overall surface charge on the cell and cocaine is an agent that eliminates a subset thereof. This description is general. Accordingly, the model can make two predictions: (i) similar currents are expected to arise upon application of any ligand to cells expressing cognate membrane protein. (ii) Similarly, inhibitors other than cocaine are expected to induce currents when applied to cells expressing SERT. We tested both predictions.

First, we challenged HEK293 cells expressing the dopamine transporter with cocaine: this gave rise to currents, which had a current-voltage relation with a negative slope (supplemental figure 1). To test the second prediction we used desipramine binding to SERT. Consistent, with the results obtained with cocaine we detected an inwardly directed peak current upon rapid application of 100 μM desipramine (Fig. 8a). We also recorded a drop in apparent membrane capacitance, if cells expressing SERT were challenged with 10 μM desipramine (representative trace in the left panel of Fig. 8b). This was not seen in control cells (right panel of Fig. 8b). At concentrations exceeding 10 μM, desipramine increased membrane capacitance in both, SERT expressing and control cells (representative traces in Fig. 8c). Thus, in SERT expressing cells, the concentration-response curve for the desipramine-induced change in capacitance was biphasic with a decline in the low concentration range followed by an increase in the high concentration range (full symbols in Fig. 8d). In contrast, only the ascending limb of the concentration-response curve was seen in untransfected control cells (empty symbols in Fig. 8d).

**Figure 8.**
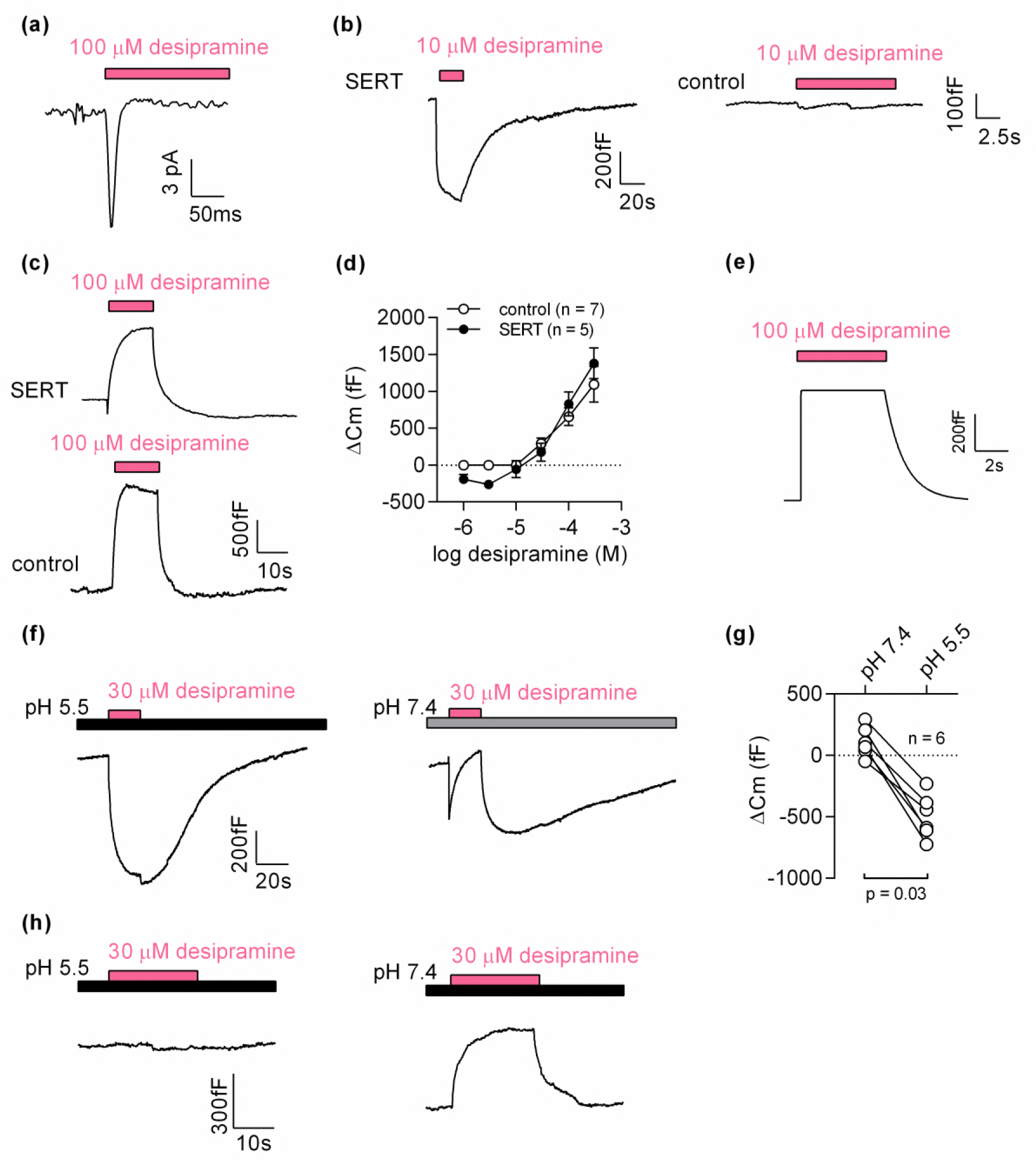
Desipramine neutralizes surface charges on SERT at low concentrations and changes the surface charge density at the inner leaflet of the membrane at high concentrations. (a) Representative displacement current evoked by the application of 100 μM desipramine to a SERT expressing cell. (b) Application of 10 μM desipramine resulted in a reduction of membrane capacitance in a cell overexpressing SERT (left panel) but not in untransfected control cell (right panel). (c) Application of 100 μM desipramine gave rise to an apparent increase in membrane capacitance in both, SERT expressing cells and control cells. (d) Concentration-dependence of the desipramine induced change in capacitance (ΔCm) in SERT expressing cells and in control cells. Data are means ± SD (control: n = 7, SERT: n = 5). (e) Simulated change in capacitance following charge elimination on the intracellular membrane surface. (f) Representative capacitance change in SERT expressing cells in response to 30 μM desipramine at pH 5.5 (left panel)) and at pH 7.4 (right panel). (g) Comparison of ACm upon application of 30 μM desipramine at pH 7.4 and pH 5.5. The lines connect data recorded in the same cell: pH 7.4: 111 ± 122 fF; pH 5.5: −496 ± 179 fF; n=6; p=0.03, Wilcoxon test. (h) Representative capacitance change in response to 30 μM desipramine at pH 5.5 (left panel) and at pH 7.4 (right panel) in a control cell. Paired measurements in 7 control cells showed no change upon application of 30 μM desipramine at pH 5.5, and a change of 332 ± 68 fF at pH 7.4 (data not shown).

### Desipramine binding to the cytosolic surface increases the membrane capacitance

Due to its tricyclic dibenzazepine structure, desipramine is very lipophilic. The increase in capacitance, which was seen at desipramine concentrations **≥** 30 μM in both SERT expressing and untransfected control HEK293, can be rationalized by assuming that desipramine permeated the cell membrane and adsorbed to the inner surface. We carried out a simulation based on this conjecture, which recapitulated the capacitance recordings (Fig. 8e). Thus, the rise in capacitance was the expected consequence of desipramine adsorption to the inner surface of the membrane. We verified this conclusion by lowering the pH in the bath solution to 5.5. This manipulation increases the fraction of protonated species of desipramine at the expense of the diffusible uncharged species (pKa =10.2). Lowering the pH is therefore predicted to diminish the SERT-independent increase in membrane capacitance, if this action is contingent on the presence of intracellular desipramine. This was the case: at pH 5.5, 30 μM desipramine caused a decrease in apparent membrane capacitance (left panel in Fig. 8f). In contrast, at pH 7.4, 30 μM desipramine elicited an increase in capacitance (right panel in Fig. 8f). It is also evident from this representative trace (right panel in Fig. 8f) that the increase at pH 7.4 was preceded by an initial decrease. This can be explained as follows: the initial decrease is the consequence of desipramine binding to SERT; the subsequent increase results from accumulation of intracellular desipramine. When assessed in paired recordings at pH 7.4 and pH 5.5, 30 μM desipramine consistently produced a drop in capacitance at pH 5.5 but not at pH 7.4 (Fig. 8g). We also repeated these experiments in control cells (Fig. 8h). As expected desipramine increased the capacitance at pH 7.4 (ΔC ∼300 fC) but was ineffective at pH 5.5.

### Ibogaine binds to an extracellular site on SERT

Most inhibitors of SERT bind to the outward facing conformation. However, ibogaine is one notable exception. Ibogaine stabilizes the inward facing conformation of SERT and inhibits substrate uptake in a non-competitive fashion (Jacobs et al., 2007). Ibogaine was therefore assumed to act on SERT by binding to a site in the inner vestibule. This implies that ibogaine gains access to the inner vestibule through the cytosol. We examined this hypothesis by measuring the effect of ibogaine on membrane capacitance (Fig. 9): application of 10 μM ibogaine, resulted in a reduction of the membrane capacitance (upper panel in Fig. 9a). This effect was absent in control cells (lower panel in Fig. 9a). Hence, these observations unequivocally refute the idea that ibogaine binds to a cytosolic site on SERT. Instead, our data suggest that the ibogaine binding site is directly accessible from the extracellular face of the transporter. We also evaluated the ibogaine-induced change in membrane capacitance in paired recordings at pH 5.5 and 7.4, which did not affect the response to ibogaine (figure 9b), although the difference in pH is expected to substantially decrease the fraction of non-protonated, membrane permeable ibogaine (pKa = 8.05).

**Figure 9.**
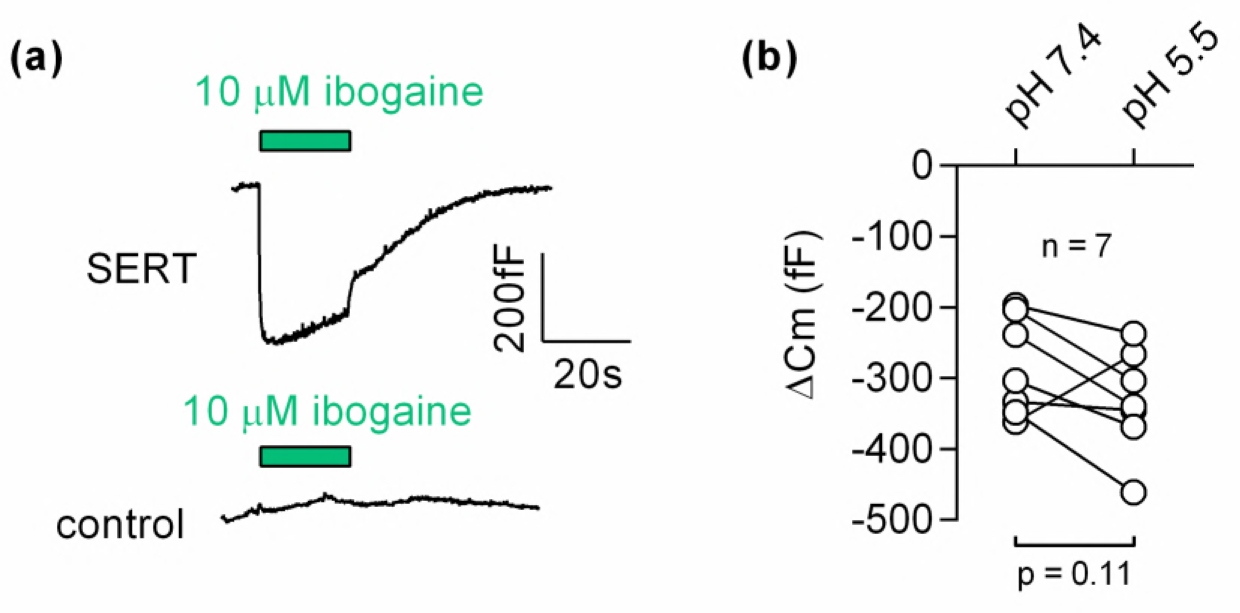
Ibogaine binds to an extracellular site of SERT. (a) Representative capacitance change induced by 10 μM ibogaine in a SERT expressing cell (upper panel) and a recording in a control cell (lower panel) (b) A comparison of the ibogaine-induced reduction in membrane capacitance at pH 7.2 and 5.5. The lines connect data points measured in the same cell (n=7). The capacitance changes were 283 ± 70 fF and 331 ± 73 fF at pH 7.4 and pH 5.5, respectively (means ± S.D.). These values were not significantly different (p=0.11; Wilcoxon test)

### Discussion

The Gouy-Chapman model was developed more than a century ago to understand the distribution of diffusible ions residing on top of a charged and polarizable surface (Gouy, 1909; Chapman, 1913). While originally developed for a description of the interaction between dissolved ions and a metal electrode, this model also provides a conceptual framework to approach the electrical properties of a lipid bilayer. Fixed charges arise primarily from the net negative charge of phospholipid head groups (McLaughlin et al., 1971). In addition, proteins, which reside in or at the plasma membrane, also contribute to the surface charges (Green and Andersen, 1991; Madeja, 2000). In cells, these surface charges are asymmetrically distributed with the inner membrane leaflet carrying a higher negative surface charge density. The electrostatic forces between the dissolved ions and the fixed surface charges of the membrane drives the formation of a Gouy-Chapman diffusive layer. Thus, the underlying theory predicts, under voltage clamp conditions, any binding reaction, which eliminates or adds a charge to the surface of the cell membrane must result in a detectable displacement current and an apparent change in capacitance. In spite of the veneration of the Gouy-Chapman model, the current experiments are – to the best of our knowledge - the first to demonstrate that binding reactions can be directly monitored by recording the predicted currents and the apparent changes of the membrane capacitance.

In the present study, we exploited the rich pharmacology of SERT to test the predictions of the Gouy-Chapman model: (i) Inhibitors such as cocaine and desipramine produced a displacement current and the related change in apparent membrane capacitance. Their binding *per se* did not result in movement of mobile charges within the membrane electric field and was fully accounted for by the neutralization of a surface charge. This conclusion is supported by the negative slope of the current-voltage relation. In addition, the change in membrane capacitance is consistent with the average density of SERT in the cell line employed: neutralization of one solvent accessible (=surface) charge/molecule gives rise to a change in membrane capacitance of about 500 fF. (ii) Displacement currents were also seen with the substrate serotonin provided that chloride was omitted to eliminate the conformational change associated with substrate translocation. (iii) Finally, we used ibogaine to interrogate the Gouy-Chapman model; binding of ibogaine also produced a drop in apparent membrane capacitance. While inhibitors bind to the outward facing conformation, ibogaine traps SERT in the inward facing conformation (Jacobs et al., 2007; Bulling et al., 2012). The precise binding site of ibogaine is unknown, but our capacitance recordings unequivocally refute the original hypothesis that ibogaine gains access to the inner vestibule. Our findings are also consistent with earlier observations, which showed that ibogaine was ineffective when administered to SERT via the patch electrode (Bulling et al., 2012).

SERT ligands are protonated amines, which form an ion pair with D98 in the orthosteric S1 binding site (Coleman et al., 2016) and thus neutralize a surface charge. Because of the large number of transporters expressed on the surface of our HEK293 cell line, neutralization of one aspartate per transporter suffices to account for our experimental findings, both qualitatively and quantitatively. However, it is plausible that the binding of an uncharged ligand can also be detected by a capacitance change provided that ligand binding results in conformational rearrangements, which alter the solvent accessible surface of the protein. We note that on average the changes in apparent capacitance were somewhat larger in with cocaine (495 fF, n=29) than with ibogaine (283 fF, n=7; p=0.0007, Mann-Whitney U-test). This difference presumably reflects the difference in accessible surface charges in the cocaine- and the ibogaine-bound state. Finally, it is worth pointing out that capacitance are more specific for monitoring binding events than the recording of the displacement current. The displacement currents evoked by cocaine, desipramine and 5-HT (in the absence of Cl^-^) validate the predictions of the Gouy-Chapman model. However, the small size and transient occurrence of these currents limit their applicability for the assessment of ligand/protein interactions. Recordings of the apparent change in membrane capacitance is a better technique to investigate ligand binding because of the sustained amplitude change (figure 6). An additional advantage of this readout is that measurements of membrane capacitance are less susceptible to the masking effects of electrogenic transitions associated with conformational change. This was also seen in our recordings: the 5-HT induced peak current was approximately 4 times larger than the cocaine peak, because most of the current elicited by 5-HT was produced by charges moving in response to conformational change. Thus, despite the small valence (0.15) associated with this 5-HT-induced conformational rearrangement the displacement current was masked beyond recognition. In contrast, the extent by which the capacitance was reduced was similar for cocaine and 5-HT (see figure 6f). This can be explained as follows: in a cell expressing 20 million SERT molecules, a 5-HT-induced conformational rearrangement associated with a valence of 0.15 can at maximum change the total membrane capacitance by about 280 fF. This is less than the 500 fF change, which is predicted to result from the elimination of one charge on the outer surface of the same number of transporters. In this context we want to note that we estimated the minimal expression density of the target membrane protein required for detection of the ligand-induced change in apparent membrane capacitance to be 20,000 units/pF (see supplement, section 4).

We stress that elimination of surface charge by ligands does not evoke an actual change in membrane capacitance. Accordingly, we refer to the recorded alterations as changes in “apparent capacitance”. The point is illustrated by the equivalent circuit of a cell (figure 10): the schematic representation depicts a battery (VΦ), which is connected in series with a capacitor (C_m_) representing the cell membrane. This battery was included to account for the voltage drop caused by the asymmetry of the surface charge densities at the outer and inner membrane leaflets. Importantly, this battery is the element in the circuit, which is affected by the ligand-induced elimination of surface charges. The physiological relevance of this battery in a living organism is readily appreciated: it is, for instance, known that the voltage for half-maximal activation (V_0.5_) of voltage-gated ion channels can be shifted by the presence of divalent ions in the extracellular fluid (Frankenhaeuser and Hodgkin, 1957). This is the reason why hypocalcemia causes seizures. The shift in V_0.5_ is accounted for by the screening of surface charges by dissolved ions, which - similar to the ligand-induced elimination of surface charge -impinges on the highlighted battery. However, while surface charge screening is a long-range effect of ions that keep their hydration shell, surface charge neutralization/elimination reflects a more specific interaction: it relies on tight binding of a ligand, which requires partial shedding of its water shell. We stress that in the case of surface charge screening no reference is made to the concept of binding affinities, which -in contrast - is implicit to ligand-induced surface charge neutralization/elimination.

**Figure 10.**
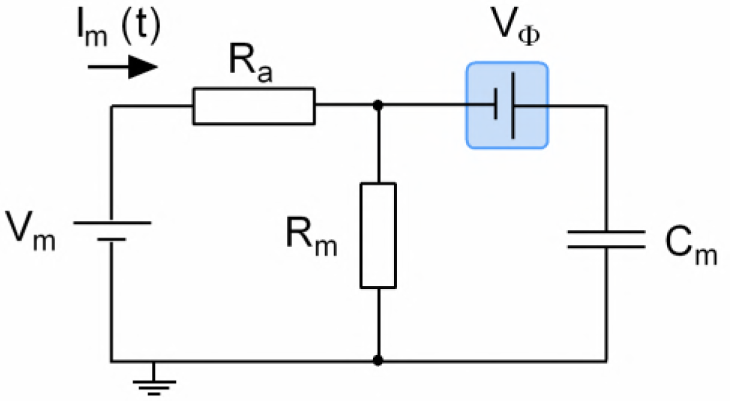
Equivalent circuit of the cell. The battery denoted as V_m_ accounts for the difference in potential between the intra- and extracellular bulk solution. R_a_ is the access resistance of the patch electrode, R_m_ is the electrical resistance of the membrane and C_m_ is the electrical capacitance thereof. Highlighted in blue is a second battery denoted by V_Φ_. This battery accounts for the potential difference created by the asymmetry in the intra- and extracellular surface charge densities and is the element in the circuit, which is affected by ligand-induced elimination of surface charge.

As a mean to study the kinetics of ligand binding, capacitance recordings are not *a priori* limited to charged ligands. In fact, any reaction, which alters the surface charge in the membrane can be detected by recording the change in apparent capacitance. This conclusion is also supported by our observations that desipramine increased the apparent membrane capacitance at concentrations exceeding 10 μM. Three lines of evidence indicate that this was a non-specific action: (i) the increase in capacitance was seen regardless of whether cells expressed SERT or not; (ii) it was not saturable and (iii) it was not mimicked by the other SERT ligands (cocaine, serotonin and ibogaine). The most plausible explanation is to assume enrichment of desipramine at the inner leaflet of the membrane. The polar surface area is a good predictor of membrane permeability (Palm et al., 1998). The polar surface areas for desipramine, cocaine and 5-HT are 15 Å^2^, 56 Å^2^ and 62 Å^2^ respectively (values taken from https://www.ncbi.nlm.nih.gov/pccompound). Thus, the low polar surface area of desipramine is consistent with its rapid permeation and the resulting change in capacitance. Incidentally, these observations highlight applications of possible membrane capacitance recordings, which go beyond measuring ligand binding in real time: there are many biological processes, which are known to alter the amount and the distribution of solvent accessible charge in the membrane. This (non-exhaustive) list includes: (i) reactions, which are orchestrated by flippases and floppases and which maintain the vital asymmetry between inner and outer surface-charge densities (Zachowski et al., 1989; Groen et al., 2011); (ii) dissipation of an existing charge-asymmetry by scramblases (Zachowski, 1993; Suzuki et al., 2010, 2013); (iii) alteration of the charge density at the inner surface by enzymes, e.g. phospholipase C via cleavage of PIP_2_ (Maucó et al., 1979; Rittenhouse-Simmons, 1979), lipid kinases (Pike and Arndt, 1988) and phosphatases (i.e. P_ten_, Maehama & Dixon 1998). These activities ought to be amenable to recordings of the membrane capacitance given that they result in a sufficiently large change in surface charge density. Thus, capacitance measurements may be useful to extract kinetic information on these key biological reactions in real time and by a label-free approach.

## Experimental procedures

### Whole-Cell patch clamp recordings

Recordings were performed on tetracycline-inducible HEK293 cells stably expressing the human serotonin transporter (hSERT). The cells were maintained in Dulbecco’s Modified Eagle’s Medium (DMEM) containing 10 % fetal bovine serum and selection antibiotics (150 ug-ml^-1^ zeocin and 6 ug-ml^-1^ blasticidin). Twenty-four h prior to the experiment, the cells were seeded onto poly-D-lysine-coated dishes (35 mm- Nunc Cell-culture dishes, Thermoscientific, USA) containing medium supplemented with 1 ug-ml^-1^ tetracycline. If not stated otherwise the cells were continuously superfused with an external solution containing 140 mM NaCl, 3 mM KCl, 2.5 mM CaCl_2_, 2 mM MgCl_2_, 20 mM glucose, and 10 mM HEPES (pH adjusted to 7.4 with NaOH). In some instances the following changes were made: (i) for experiments at pH 5.5, HEPES was substituted by 2-(N-morpholino)ethansulfonic-acid; (ii) for experiments in Na^+^-free external solution NaCl was replaced by NMDG^+^(N-methyl-D-glucamine)Cl-; (iii) for experiments in Cl-free external solution NaCl was replaced by Na^+^MES (methanesulfonate). The internal solution in the patch pipette contained 152 mM NaCl, 1 mM CaCl_2_, 0.7 mM MgCl_2_, 10 mM HEPES, 10 mM EGTA (ethylenglycol-bis(aminoethylether)-N,N,N’,N’-tetra-acidic-acid) (pH 7.2 adjusted with NaOH). Where indicated, the internal Cl-concentration was reduced (152 mM NaOH, 1 mM CaCl_2_, 0.7 mM MgCl_2_, 10 mM EGTA, 10mM HEPES, pH 7.2 adjusted with methaneosulfonic acid). Ligands were applied via a 4 tube or 8 tube ALA perfusion manifold using the Octaflow perfusion system (ALA Scientific Instruments, USA), which allowed for complete solution exchange around the cells within 100 ms. Currents were recorded at room temperature (20-24 °C) using an Axopatch 200B amplifier and pClamp 10.2 software (MDS Analytical Technologies). Current traces were filtered at 1 kHz and digitized at 10 kHz using a Digidata 1440 (MDS Analytical Technologies). The recordings were analyzed with Clampfit 10.2 software. Passive holding currents were subtracted, and the traces were additionally filtered using a 100-Hz digital Gaussian low-pass filter.

### Membrane capacitance measurements

Recordings were performed in the whole-cell configuration. For the measurement we used a train of square wave voltage pulses with an amplitude ± 40 mV and a frequency of 200 Hz. The holding potential was set to 0 mV if not stated otherwise. Exponential current responses were low pass filtered by a 10 kHz Bessel filter and sampled at 100 kHz rate. The acquired current traces were first deconvoluted with the transfer function of the recording apparatus and the passive membrane parameters of a cell were calculated from the theoretical function as described by Hotka & Zahradník (2017). The pipette capacitance was recorded in the cell-attached mode first and subtracted from the currents recorded in the whole-cell configuration prior to further analysis. The patch pipettes used had a resistance of 2-4 MΩ. To stabilize the level of stray capacitance, the pipettes were coated with hydrophobic resin Sylgard184 (Dow Corning, USA).

## Authors Contributions

W.S., M.H. and V.B designed experiments; Y.L. and V.B. recorded displacement currents; V.B. conducted the capacitance measurements; V.B. analyzed the data; M.H. and W.S. modeled the data; V.B, M.H., M.F. and W.S. interpreted the data and wrote the paper.

## Acknowledgements

We thank Shreyas Bhat, Klaus Schicker, and Peter S Hasenhuetl for discussions and comments on the data.

## Footnotes

This work was supported by the Austrian Science Fund/FWF Project P28090 (to W. S.), Project Program Grant SFB35 (F3510 to M. F.). The authors declare that they have no conflicts of interest with the contents of this article.

